# Engineered Antiviral Sensor Targets Infected Mosquitoes

**DOI:** 10.1101/2023.01.27.525922

**Authors:** Elena Dalla Benetta, Adam J. López-Denman, Hsing-Han Li, Reem A. Masri, Daniel J. Brogan, Michelle Bui, Ting Yang, Ming Li, Michael Dunn, Melissa J. Klein, Sarah Jackson, Kyle Catalan, Kim R. Blasdell, Priscilla Tng, Igor Antoshechkin, Luke S. Alphey, Prasad N. Paradkar, Omar S. Akbari

## Abstract

Escalating vector disease burdens pose significant global health risks, so innovative tools for targeting mosquitoes are critical. We engineered an antiviral strategy termed REAPER (v<u>**R**</u>NA <u>**E**</u>xpression <u>**A**</u>ctivates <u>**P**</u>oisonous <u>**E**</u>ffector <u>**R**</u>ibonuclease) that leverages the programmable RNA-targeting capabilities of CRISPR Cas13 and its potent collateral activity. Akin to a stealthy Trojan Horse hiding in stealth awaiting the presence of its enemy, REAPER remains concealed within the mosquito until an infectious blood meal is up taken. Upon target viral RNA infection, REAPER activates, triggering programmed destruction of its target arbovirus such as chikungunya. Consequently, Cas13 mediated RNA targeting significantly reduces viral replication and its promiscuous collateral activity can even kill infected mosquitoes. This innovative REAPER technology adds to an arsenal of effective molecular genetic tools to combat mosquito virus transmission.

**One-Sentence Summary:** Engineered Cas13-based antiviral sensor kills infected mosquitoes upon activation from arbovirus infection.

## Introduction

Arboviruses are among the most widespread pathogens affecting humans, causing deadly diseases worldwide, and their geographic distribution and incidences are escalating globally due to a changing climate. RNA arboviruses belong to the families *Flaviviridae, Togaviridae*, and *Bunyaviridae* and are primarily transmitted by mosquitoes ^1^. More specifically, *Aedes* mosquitoes notoriously transmit numerous arboviruses, such as chikungunya (CHIKV), dengue (DENV), yellow fever (YFV), and Zika (ZIKV), that together infect hundreds of millions of people annually. Existing traditional vector control strategies have not adequately curtailed either virus transmission nor the spread of mosquitoes to new habitats ^2^. Therefore, novel vector-control strategies are required.

Historically, efforts to reduce viral transmission have focused on the use of chemical insecticides and the development of vaccines. However, with the rise of insecticide resistance ^3^ and the limited efficacy of vaccines ^4^, there is a critical need for improved control strategies. Recently, molecular tools, such as broadly neutralizing antibodies ^5^ and encoded miRNA arrays ^6,7^ have been demonstrated to mitigate DENV and ZIKV transmission by mosquitoes. Additionally, CRISPR-Cas technologies have been engineered to control mosquito populations using the precision-guided Sterile Insect Technique (pgSIT) ^8^ and confinable gene drives ^9,10^. These molecular tools present great promise as vector control strategies capable of reducing viral spread by mosquitoes. However, the rapid evolution of pathogens requires the development of easily programmable technologies with direct activity against evolving and emerging infectious diseases.

Unlike traditional CRISPR technologies for insect control that target DNA and thus the mosquito vector, there are alternative systems that target RNA that could instead be effective against the arboviruses themselves. Cas13 is one such system that originally evolved as an RNA antiviral in prokaryotes ^11^, and has surfaced as programmable RNA-targeting ribonucleases ^11^. The high efficiency of programmable Cas13 has been demonstrated in human cells ^12^, zebrafish ^13^, mice ^14^, flies ^15,16^ and plants ^17^, but the high on-target activity is natively coupled with collateral cleavage of bystander RNAs ^18^. Although this collateral cleavage may limit the potential of Cas13 for therapeutic applications, we hypothesized that it could be leveraged in mosquitoes to combat viral infections, providing a basis for the development of flexible antiviral technologies.

Here we exploit the properties of Cas13 for programmable viral RNA targeting in mosquitoes. To develop this innovation, we use the *Ruminococcus flavefaciens* RfxCas13d ribonuclease (herein referred to as CasRx)^12^, which possesses robust RNA cleavage activity. We genetically encoded CasRx expression in mosquitoes and demonstrated its efficacy by targeting two non-essential genes. Subsequently, we biochemically screen the activity of gRNAs targeting CHIKV using a sensitive enzymatic nucleic acid sequence reporter (SENSR)^19^. We then use these effective gRNAs to engineer an antiviral sensor in *Aedes aegypti* termed vRNA Expression Activates Poisonous Effector Ribonuclease (REAPER). Akin to a Trojan Horse resting in wait to eliminate its enemy, REAPER remains inactive in the absence of its target viral RNA and becomes activated upon viral infection. We demonstrate that REAPER is programmed to directly target deadly arboviruses like CHIKV, resulting in reduced viral transmission and even mortality of infected mosquitoes.

## Results

### Generation of a programmable RNA-targeting platform in *Ae. aegypti*

To generate a platform for programmable RNA targeting, we created 15 transgenic *Ae. aegypti* lines that genetically encoded CasRx. CasRx expression was driven by a broadly expressed ubiquitin L40 (Ub) promoter ^20^ with either a nuclear localization signal (NLS) or a codon-optimized CasRx fused with a nuclear export signal (NES) **(Fig. S1)**. CasRx expression levels were assessed through RNA sequencing and qPCR. Lines with the highest expression levels (**Table S1, S2**) were genetically crossed with two transgenic lines expressing gRNA arrays engineered to target either an endogenous *Ae. aegypti yellow* transcript or the reporter EGFP transcript, allowing for visual quantification of expression levels. The expression of these gRNA arrays was driven by a U6 non-coding small nuclear RNA (snRNA) polymerase III promoter (U6b:gRNA_array_) ^21^ (**Fig. S1**). The resulting progeny were assessed for hatching rates, visual phenotypes, zygosity, and misexpression of the target/non-target genes quantified using both qPCR and RNA sequencing **(Fig. 1, Fig. 2, Fig. S2, Fig. S3, Fig. S4)**.

**Fig. 1.**
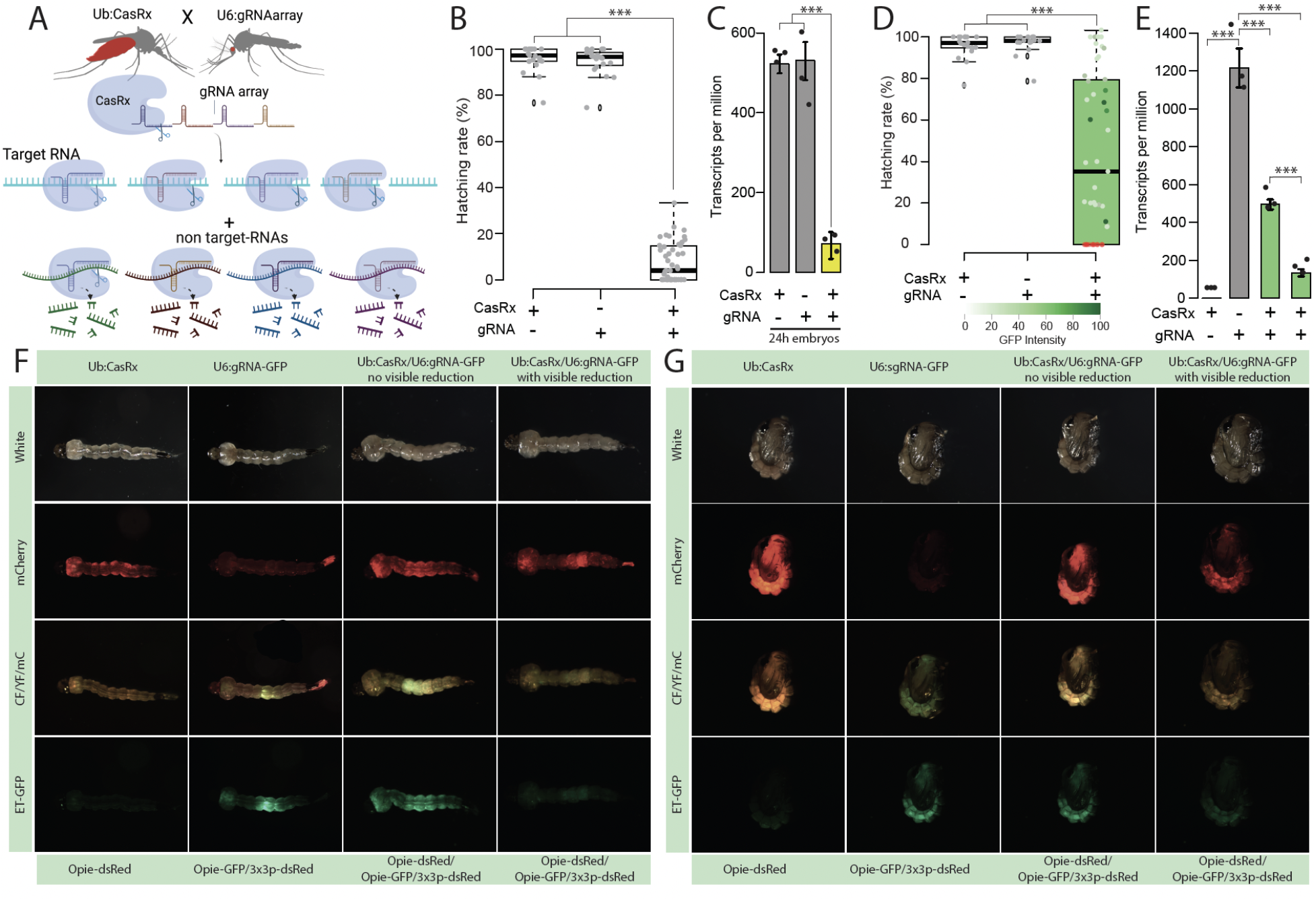
Assessment of transcript reduction mediated by CasRx. **(A)** Schematic representation of the binary system used for the proof-of-concept studies evaluating the candidate target genes. The lines are composed of CasRx and a gRNA array targeting the gene of interest inducing collateral cleavage of bystander RNAs. (**B**) Hatching rate of parental lines and transheterozygotes targeting *yellow*. **(C)** Relative expression measured by qPCR in control and transheterozygotes embryos **(D)** Hatching rate of parental lines and transheterozygotes targeting EGFP. Phenotype penetrance is depicted by green shading in the box plot, with colors ranging from light green (low EGFP levels) to dark green (high EGFP levels). Red dots represent dead individuals and thus non quantifiable EGFP. Asterisks indicate a significant reduction in the hatching rate in transheterozygotes by one-way ANOVA with Tukey’s multiple-comparison test (*** *P* < 0.001). (**E**) Relative expression measured by qPCR in control and transheterozygote larvae in EGFP. The two transheterozygote groups represent larvae with no visible reduction in EGFP and with visible reduction of EGFP, respectively. asterisks indicate significant differences by one-way ANOVA with Tukey’s multiple-comparison test (\*\*\**P* < 0.001), (**F and G**) EGFP reduction at larval and pupal stages, respectively. mCherry represents the red fluorescent filter, ET-EGFP represents the green fluorescent filter, and CF/YF/mC represents the triple hybrid filter for detecting blue, yellow, and red fluorescence.

**Fig. 2.**
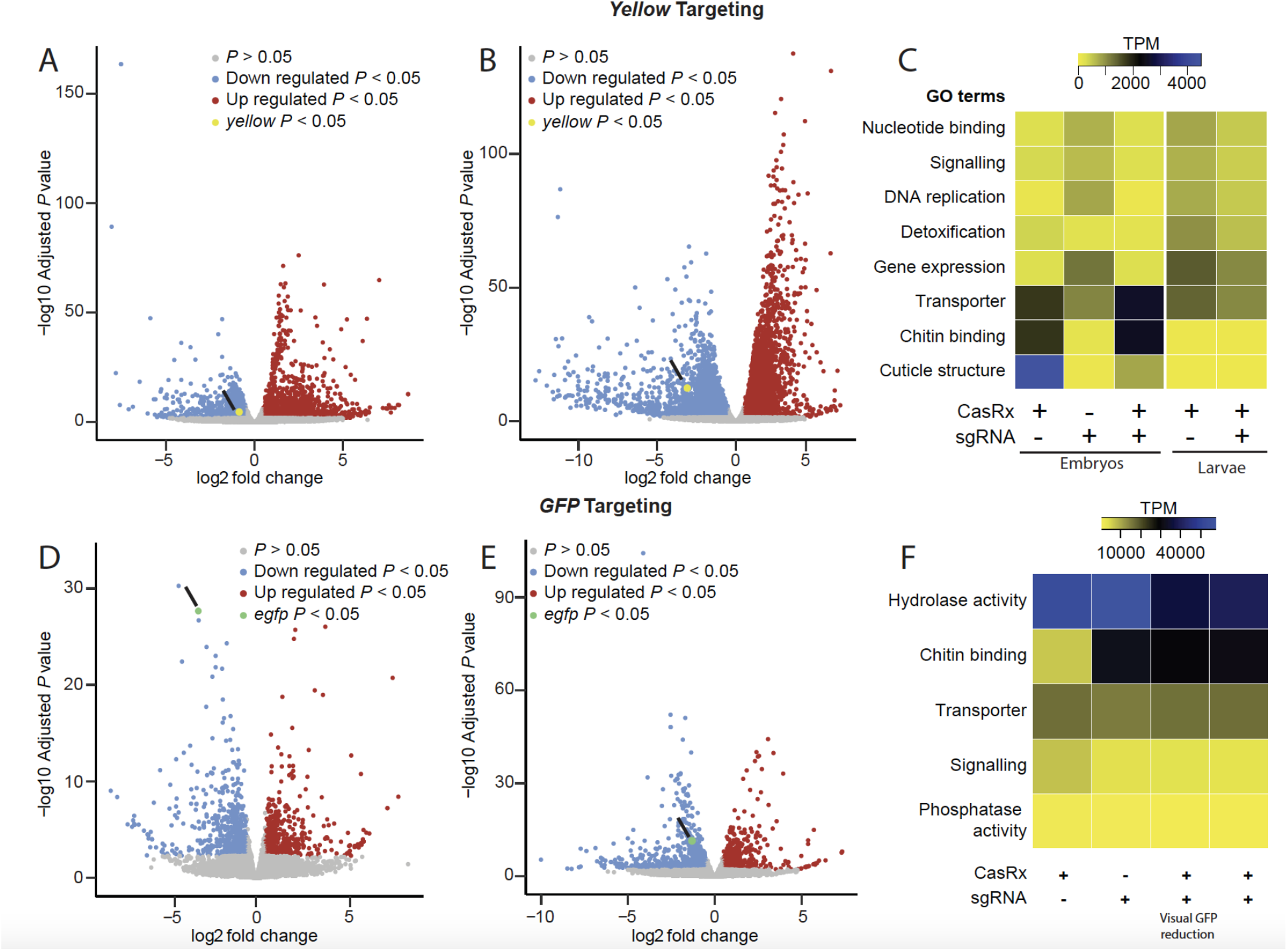
Transcriptome differential expression analysis. **(A)** RNAseq results in 24–hr-old eggs and (**B)** in third instar larvae comparing transheterozygotes targeting *yellow* to the CasRx expressing control line. Blue dots represent downregulated genes with *P* < 0.05, and red dots represent overexpressed genes with *P* < 0.05. (**C)** Heat map of gene functions with the two-fold difference in expression level among different treatments and FDR < 0.05 when targeting the *yellow* gene. Expression levels are represented by colors ranging from yellow (lowest expression) to blue (highest expression) (**D)** RNAseq results in third instar larvae with a visible EGFP reduction and (**E)** in the third instar larvae that do not show visible EGFP reduction comparing transheterozygotes targeting *GFP* to CasRx expressing control line. Blue dots represent downregulated genes with *P* < 0.05 and red dots represent overexpressed genes with *P* < 0.05. (**F)** Heat map of gene functions with the two-fold difference in expression level among different treatments and FDR < 0.05 when targeting EGFP transgene. Expression levels are represented by colors ranging from yellow (lowest expression) to blue (highest expression).

Targeting *yellow* resulted in high embryonic lethality (**Table S3**) with only 60% of the crosses producing viable transheterozygous eggs (Ub:CasRx/+; U6b:gRNA ^yellow^/+) **(Fig. 1B, Fig. S3A)**. Remarkably, only 7.59% of transheterozygotes survived to adulthood, despite *yellow* being dispensable for survival ^22^. Both qPCR and RNA sequencing detected a remarkable 13-fold reduction in *yellow* transcripts in the transheterozygote embryos as compared to controls, confirming programmed on-target disruption (**Fig. 1C; Fig. S2A**). Targeting EGFP also resulted in lethality and a reduced hatching rate of ~43% (**Fig. 1D**). The surviving transheterozygotes were evaluated by visually quantifying EGFP fluorescence (**Fig. 1F, G**), which was reduced by an average of 70.27% (**Fig. 1D, Table S5**). Both qPCR and RNAseq analysis revealed a significant 2.11–4.36-fold reduction of EGFP in larval samples (**Fig. 1E, Fig. S2B, Table S6**), indicating that even though visible GFP protein reduction was difficult to score in some surviving larvae, GFP mRNA was indeed reduced **(Fig. 1E)**. Taken together, these findings provide compelling evidence of the programmable ability of CasRx to reduce the expression of target genes, though the high mortality rates suggest toxicity, presumably due to the known collateral activity of CasRx ^15^.

To estimate the degree of CasRx collateral activity, we performed differential expression using transcriptome wide RNA sequencing (**Fig. 2, Table S6–S17**). Comparing transheterozygotes targeting *yellow* at the embryonic stage to the controls showed that remarkably ~45% of the transcriptome was misexpressed (FDR < 0.05) (**Fig. 2A, C, Table S11, S12**). The main category of misexpressed genes consisted of genes involved in structural components of the cuticle (fold change > −10; FDR < 0.05), which perhaps is indicative of collateral activity largely occurring in cells that express *yellow*, as this is an important factor in cuticle pigmentation (**Fig. 2C**, **Table S13**). In surviving transheterozygotes, only 24% of the genes were misexpressed, though these also had much lower fold changes (**Fig 2B**, **C**, **Table S14**). Comparing the differential expression between transheterozygotes targeting EGFP and the controls showed that ~10% of the transcriptome was misexpressed. Half of these misexpressed genes, representing ~5% of the transcriptome, were downregulated genes involved in signaling pathways and transporter activity (**Fig. 2D, E, F, Tables S15-S17**). Similar results were also observed when using CasRx fused with a nuclear export signal (NES) (**Fig. S4, Tables S18-S19**), indicating that the presence of NLS or NES does not affect CasRx activity. These data confirm the programmable knockdown of targeted genes, though with a significant impact on the transcriptome.

### Generation of REAPER in *Ae. aegypti*

To exploit the collateral activity of CasRx, gRNAs targeting CHIKV were synthesized and screened for their *in vitro* collateral cleavage efficiency. Four CHIKV coding regions of the non-structural protein (nsp) were targeted by designing two gRNA targeting nsp1 and nsp2 regions, and two targeting the nsp4 coding region for screening (**Table S20, File S1, Fig. S5**). To determine which gRNA resulted in the highest level of collateral activity, we used SENSR ^19^ in which the gRNAs were independently challenged against synthetic ssRNA templates mimicking the respective virus. Here, the presence of the viral template triggers the activation of CasRx/gRNA complex activity *in vitro*, and a fluorescent readout indicates the effectiveness of the deployed gRNAs for inducing collateral activity. Upon activation, the CasRx collateral activity cleaves a quenched probe to produce a fluorescent signal (**Fig. 3A,B**). All CHIKV gRNAs resulted in rapid collateral activity (**Fig. 3C**). Importantly, the data indicate that in the absence of the target RNA, the collateral activity is null, confirming that the presence of a target sequence is necessary to activate CasRx activity ^19^ (**Fig. 3B, 3D**). We noticed here that a high level of synthetic target seemed to trigger greater collateral activity. To confirm this observation and identify how target concentration influences collateral activity, we next performed a serial dilution of the synthetic templates in the SENSR assay. The results demonstrated a direct relationship between target concentration and collateral activity (**Fig. 3D, E**), confirming that high target levels trigger high collateral activity.

**Fig. 3.**
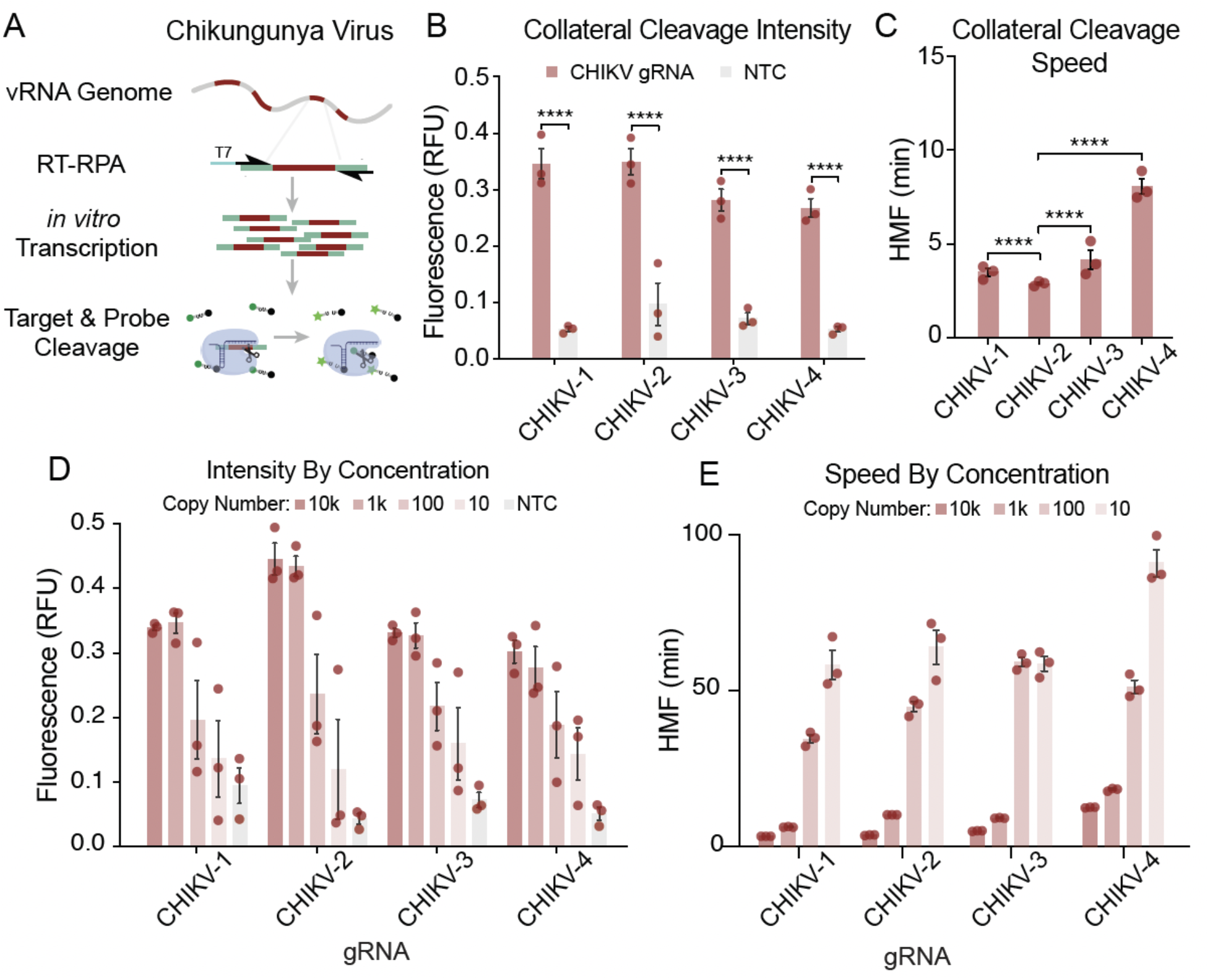
Analysis of gRNA-dependent collateral activity targeting CHIKV. **(A)** Schematic representation of the SENSR system used to determine the collateral cleavage activity of selected gRNAs. The detection protocol requires that the specific target sequences within the viral RNA are reverse transcribed (RT) into cDNA and amplified by RPA at 42 °C for 30 min. During amplification, a T7 promoter is incorporated into the 5′ terminus of the amplicons (T7, blue). In the next step, both *in vitro* transcription and CasRx collateral cleavage occur simultaneously. Finally, the recognition and cleavage of the target RNA sequence complementary to the gRNA induce collateral cleavage of bystander RNA molecules. Collateral cleavage of a modified probe conjugated to 6-FAM, and a fluorescence quencher facilitates readout by fluorescence. **(B)** Initial characterization of the collateral activity for each gRNA targeting CHIKV. SENSR analysis was performed against 10,000 copies/μL of the synthetic target. The fluorescence signal represents the background-subtracted signal. Statistical significance was calculated using Sidak’s multiple comparisons test (n = 3). **(C)** Collateral cleavage speed of each gRNA calculated using half-maximum fluorescence (HMF) analysis (see methods), where a lower HMF indicates faster collateral activity. Statistical significance was calculated using Tukey’s multiple comparison test (n = 3) **(D)** Intensity of signal from SENSR assay for each gRNA along a concentration gradient. The fluorescence signal represents the background-subtracted signal (n = 3). **(E)** Speed of cleavage along concentration gradient for each gRNA used to target CHIKV. The speed of collateral activity is represented using an HMF analysis (n = 3).

With these results in hand, we next engineered a transgenic line expressing the four gRNAs tested with SENSR (U6b:gRNA^CHIKV^) and crossed it to the CasRx lines, encoding either a NLS (Ub:CasRx-NLS) or NES (Ub:CasRx-NES-A and Ub:CasRx-NES-B). In order to activate the REAPER system, transheterozygous mosquitoes were challenged with a CHIKV infection and the analysis of viral reduction and/or mosquito viability at 14 days post infection was recorded to test the efficiency of these lines (**Fig. 4A)**. In the presence of Ub:CasRx-NLS, the mosquitos, challenged by a CHIKV infection, showed a minimal effect on the viral genome copy numbers, infection rates, and survival as compared to the controls (**Fig. S6**). In contrast, the U6b:gRNA^CHIKV^ mosquito line crossed with either Ub:CasRx-NES-A or Ub:CasRx-NES-B showed significant decreases in both survival, as the majority of the mosquitoes died within 5 days post CHIKV infection (**Fig. 4B**), and viral titers indicating that the NES is effective at targeting viruses like CHIKV that replicate in the cytoplasm (**Fig. 4C**). Taken together these data indicate that REAPER is an effective strategy to target vRNA and even kill infected mosquitoes.

**Fig. 4.**
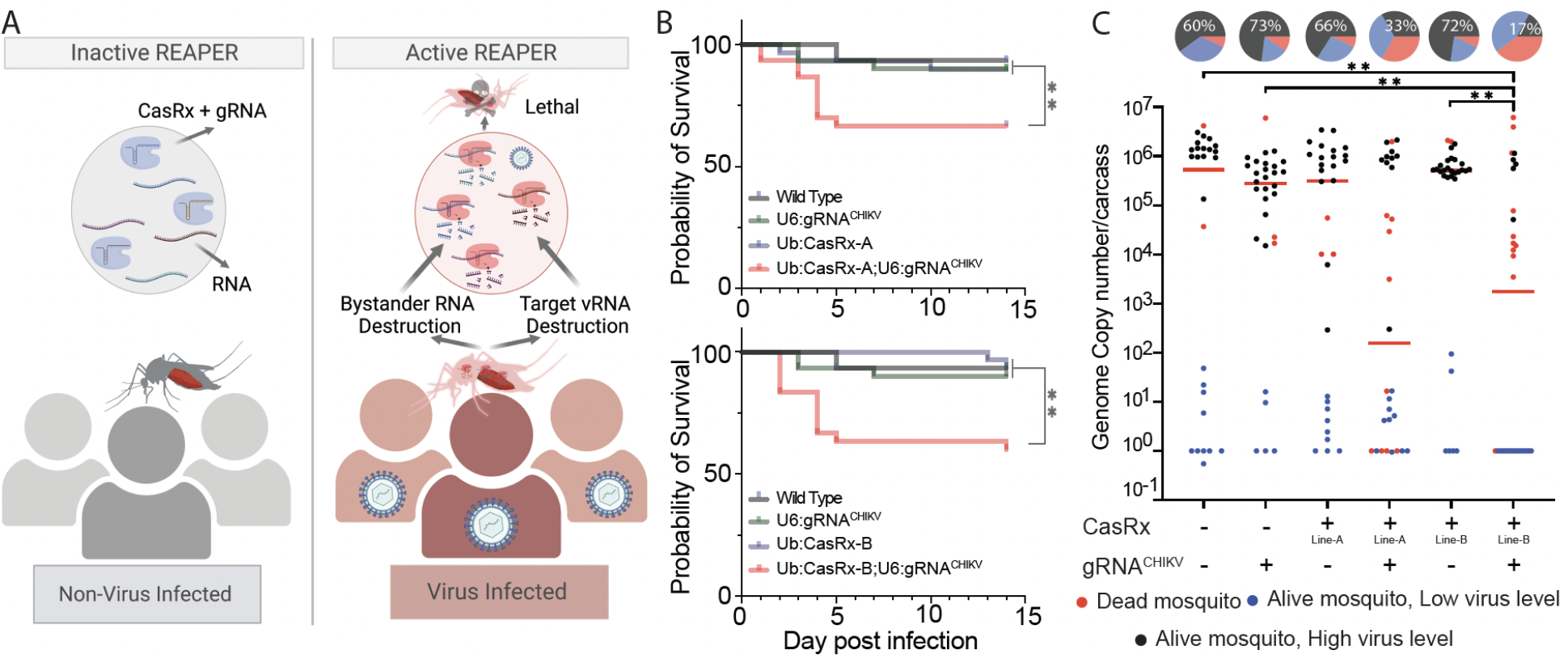
Viral challenge assays. **(A)** Schematics representation of the REAPER system. REAPER remains inactive in uninfected mosquitoes and is activated in the presence of the targeted virus. Once activated, REAPER induces degradation of viral RNA with additional collateral cleavage of bystander RNAs, leading to lethality. **(B)** Adult survival curves (log-rank Mantel-Cox test) of virus exposed females per treatment. Asterisks represent significant differences in survival between treatments(\**P* < 0.05; \*\**P* < 0.001). **(C)** The viral genome copy number and infection prevalence of CHIKV, were measured 14 days after an infected blood meal challenge (n = 30). qRT-PCR was used to assess genome copy number and infection prevalence in individual mosquitoes, with each dot representing the viral load from individual mosquitoes. Each pie-chart indicates the percentage of mosquitos that died by day 5 (in red), `that are alive but with lower virus level of ≤10`^2^ (blue), or that have an high virus level (black). Lines A and B represent different CasRx lines (**Table S2**). Horizontal red lines indicate the median of the viral loads. Considering the non-normal distribution of viral titers, the median was used to describe central tendency. The non-parametric Mann-Whitney test was used to compare median viral titers, and Fisher’s exact test was used to compare infection prevalence. \**P* < 0.05, \*\**P* < 0.01, \*\*\*\**P* < 0.0001.

## Discussion

In this study, we addressed the need for engineering programmable and effective methods for controlling viral transmission via mosquito vectors by generating a Cas13 based antiviral system termed REAPER in *Ae. aegypti*. REAPER exploits the collateral activity of CasRx to impede viral replication and/or kill infected mosquitos, limiting viral transmission. Importantly, in true Trojan horse fashion, REAPER lies in wait for the presence of its “enemy” and is only activated following viral infection. With our system, we demonstrate reduction of CHIKV replication in *Ae. aegypti*.

REAPER technology is based on efficient RNA-targeting in addition to CasRx collateral activity of bystander RNA, selecting highly efficient gRNAs is thus essential to design an effective strategy. We thus developed an approach to confirm viral targets through bioinformatic analysis and verified high collateral activity *in vitro* using the previously developed SENSR system ^19^. Rapid and robust collateral activity was observed for all gRNAs tested. Since there is a need for a technology that can detect multiple virus serotypes or different viruses, we used *in vitro* data information to develop a multiplexed strategy to target four target regions of CHIKV. Although all gRNAs tested showed this rapid and robust collateral activity, we observed differences in speed and intensity. Nevertheless all CHIKV gRNAs had fast collateral activity *in vitro* and were able to induce high lethality *in vivo*. Previous work has described differences in interference efficiencies from different gRNAs ^23,24^, suggesting that target RNA accessibility might impact CasRx activity. However, there are limited comprehensive studies of collateral activity, though a greater understanding of this phenomenon could improve Cas-based Trojan horse systems and other applications that exploit collateral activity. SENSR data confirmed that the presence of a target sequence is necessary to activate CasRx collateral activity enabling us to develop a system that is dormant in uninfected mosquitoes and activated following infection. The system reduced viral replication and therefore transmission by mosquitoes by either knocking down viral RNA expression or killing the infected mosquitoes.

The presented design of the REAPER system lays out a foundational technology that can be further optimized and expanded. In the initial designs, we compared the expression of CasRx fused with either an NLS or NES and found no considerable differences in activity against two phenotypic genes. Nonetheless, the NES is critical for targeting viruses, which replicate in the cytoplasm (**Fig. 4, Fig. S6**). This points out that the subcellular localization of a target will be an important factor to consider when designing a REAPER strategy in other organisms. We also observed transcriptome misregulation when expressing CasRx alone with a ubiquitin promoter, which may lead to toxicity, limit fitness, or reduce the efficacy of an antiviral platform. Future designs should therefore implement tissue-specific or inducible promoters for CasRx expression to reduce the deleterious effects associated with ubiquitous expression. Furthermore, gRNA design and *in vitro* screening are likely to enhance the potency of the REAPER system, exemplified through our analysis using the SENSR system (**Fig. 3**). Our results also demonstrate that multiplexed gRNAs are functional in the system (**Fig. 4**), which can permit the generation of future multi-virus antiviral systems.

Overall, our results with the REAPER system indicate that CasRx can be used to engineer interference against RNA viruses in mosquitoes. The exploitation of collateral activity is fundamental to design an effective system. Correspondingly, finding or engineering novel CRISPR ribonucleases with increased collateral activity or understanding how collateral activity is triggered by different gRNAs sequences, will be very important to improve the REAPER technology. Maximizing the speed and robustness of collateral activity will lead to the generation of novel mechanisms for RNA-guided immunity against multiple diseases in vertebrates and provides a platform to develop novel mosquito control strategies for arboviruses.

## Materials and Methods

### Mosquito rearing and maintenance

All *Ae. aegypti* lines used in this study were generated from the Liverpool strain. Colonies were reared at 27.0°C with 70–80% relative humidity and a 12-hr light/dark cycle. Adults were fed 0.3 M aqueous sucrose *ad libitum*. To collect eggs, mature females were blood-fed on anesthetized mice. Oviposition cups were provided ~3 days post blood-meal, and the eggs were collected and aged for ~4 days before hatching. Mature eggs were submerged in deionized H_2_O and placed overnight in a vacuum chamber set to 677 Millibar Pressure Unit (mbar). Emerged larvae were reared in plastic containers (Sterilite) with ~3 liters of deionized H_2_O and fed daily with fish food (Tetramin).

### Design and cloning of constructs

Gibson enzymatic assembly was used to construct all cloned plasmids. To generate the CasRx expressing construct OA-1050T (Addgene plasmid #191374), the plasmid OA-1050E (Addgene plasmid #132416) was used as a backbone. A DNA fragment containing the *Ae. aegypti* ubiquitin promoter was amplified from the Addgene plasmid #100580 using primers 874F/1050T.C1 and was inserted using the AvrII and PacI restriction enzyme sites. To generate the *Ae. aegypti* codon-optimized CasRx expressing construct OA-1163A (Addgene plasmid #194001), the OA-1050T plasmid was cut by restriction enzymes PacI and NheI to remove CasRx. The codon-optimized CasRx fragment was synthesized from GenScript (GenScript USA Inc., Piscataway, NJ) and then cloned in.

Additionally, we designed four gRNA-array constructs, OA-1085F (Addgene plasmid #191375), OA-1085L (Addgene plasmid #191376), and OA-1093B (Addgene plasmid #194003), targeting *yellow*, EGFP and CHIKV, respectively. We synthesized OA-1085F and OA-1093B using Gene Synthesis (GenScript USA Inc., Piscataway, NJ), with each plasmid containing *piggyBac*, 3xp3-tdTomato, the U6b (AAEL017774) promoter, and 3’UTR in addition to an array of four programmed transcript-targeting spacers (30 nt in length) each positioned between the CasRx-specific direct repeats (36 nt in length) with a conserved 5’-AAAAC motif at the 3′ end of the direct repeat followed by a 7-thymine terminator ^12^. To generate construct OA-1085L, a plasmid (Addgene #120363) containing *piggyBac* and 3xp3-tdTomato was used as the backbone, which was linearized at the restriction enzyme sites AvrII and AscI to clone in the designed fragment.

This insert fragment was synthesized using Gene Synthesis (GenScript USA Inc., Piscataway, NJ) to contain the U6b (AAEL017774) promoter and 3’UTR in addition to an array of four programmed GFP transcript-targeting spacers (30 nt in length) each positioned between the CasRx-specific direct repeats (36 nt in length) with a conserved 5’-AAAAC motif at the 3′ end of the direct repeat followed by an extra DR, a 7-thymine terminator and then the Opie2-EGFP marker. All primer sequences can be found in **Table S21**. All plasmids and annotated DNA sequence maps are available at www.addgene.com under accession numbers: 191374 (OA-1050T), 191375 (OA-1085F), 191376 (OA-1085L), 194001 (OA-1163A), 194003 (OA-1093B).

### Generation of transgenic lines

Transgenic lines were generated by microinjecting pre-blastoderm embryos (0.5–1 hours old) with a mixture of the piggyBac plasmid (200 ng/μl) and a transposase helper plasmid (phsp-Pbac; 200 ng/μl). Embryonic collection and microinjections were performed following previously established procedures ^25,26^. Four days post-microinjection, G_0_ embryos were hatched in deionized H_2_O under vacuum (20 inHg). Surviving G_0_ pupae were sex-separated into ♀ or ♂ cages (5 G0s per cage). WT pupae of the opposite sex and of a similar age were added to cages at a 5:1 ratio (WT:G_0_). Several days post-eclosion (~4–7), a blood-meal was provided, and the eggs were collected, aged, and hatched. Hatched G_1_ larvae were screened and sorted for the expression of relevant fluorescent markers using a fluorescent stereo microscope (Leica M165FC). To isolate separate insertion events, selected transformants were individually backcrossed to WT (5:1 ratios of WT:G0), and separate lines were established. Each individual line was maintained as mixtures of homozygotes and heterozygotes with the periodic selective elimination of WTs. In total, 5 Ub:CasRx-NLS, 3 U6b:gRNA^yellow^, 1 U6b:gRNA^EGFP^, 10 Ub:CasRx-NES, and 1 gRNA lines targeting CHIKV were created (**Table S2**).

### Generating and screening for CasRx/gRNA transheterozygote

To genetically assess the efficiency of CasRx ribonuclease activity, we bidirectionally crossed the highest Ub:CasRx expressing lines to gRNA-expressing lines targeting either the *yellow* (Ub6:gRNA^*yellow-A*^, Ub6:gRNA^*yellow-B*^, Ub6:gRNA^*yellow-C*^) or EGFP (Ub6:gRNA^*EGFP*^) transcripts. After allowing the crosses to mate for 3 days, females were blood-fed for 2 consecutive days. Three days after blood-feeding, females were individually captured in plastic vials lined with moistened paper. Captured females were kept for 2 days to allow for egg laying and were removed afterwards. Collected eggs were either processed for RNA collection or hatched to screen progeny. The number of eggs on the moistened paper and larvae were then recorded and used to calculate the hatching rate. The larvae were also screened for the number of transheterozygotes, which was used to calculate the inheritance and penetrance of the observable phenotypes. Mosquito larvae and pupae were imaged using different filters (ET-EGFP, CF/YF/mC, mCherry, white) on aLeica M165FC fluorescent stereomicroscope equipped with a Leica DMC4500 color camera. The hatching rate was calculated as the ratio of number of larvae to the number of eggs, and the penetrance of the EGFP-targeting phenotype was calculated as the ratio between the number of transheterozygotes showing EGFP reduction and the total number of transheterozygotes. Statistical analyses were performed using the one-way ANOVA and Tukey’s multiple-comparison tests in RStudio (version 1.2.5033, © 2009–2019).

### Total RNA collection and qPCR

To directly select the lines with the highest CasRx expression, 10 adult individuals were collected from each of the 5 selected Ub:CasRx-NLS lines and from each of the 10 selected Ub:*Ae*CasRx-NES lines listed in **Table S2**. Additionally, to directly observe and quantify the reduction in targeted *yellow* and EGFP transcripts by the Ub:CasRx-NLS or Ub:*Ae*CasRx-NES lines, we collected transheterozygous embryos and transheterozygous larvae for RNA extraction and subsequent quantitative PCR and RNA sequencing. Embryos were collected 24 hrs post-oviposition from the F1 transheterozygous line (Ub:CasRx-NLS/U6b-gRNA^*yellow-A*^) as well as the parental lines (Ub:CasRx-NLS; U6b-gRNA^*yellow-A*^). Three biological replicates were collected per line for a total of 15 samples.

Additionally, to quantify the knockdown of *yellow* and EGFP in surviving larvae, third instar larvae were used for RNA extraction and subsequent qPCR analysis and RNA sequencing. For this, another three biological replicates were collected per line for a total of 18 samples, including 3 transheterozygous lines (1 targeting yellow and 2 targeting EGFP at different visual levels) and the 3 parental controls. Embryos and larvae were also collected for qPCR analysis of Ub:*Ae*CasRx-NES crosses with gRNA lines targeting *yellow* and EGFP. Total RNA was extracted using a Qiagen RNeasy Mini Kit (Qiagen 74104). Following extraction, the total RNA was treated with an Invitrogen DNase treatment kit (Invitrogen AM1906). RNA concentration was analyzed using a Nanodrop OneC UV-vis spectrophotometer (ThermoFisher NDONEC-W). About 1 μg of total RNA was used to synthesize cDNA with a RevertAid H Minus First Strand cDNA Synthesis kit (Thermo Scientific). cDNA was diluted 50 times before use in Real-Time quantitative PCR (RT-qPCR). RT-qPCR was performed with SYBR green (qPCRBIO SyGreen Blue Mix Separate-ROX Cat #: 17-507B, Genesee Scientific), using 4 μl of diluted cDNA for each 20 μl reaction containing a final primer concentration of 200 nM and 10 μl of SYBR green buffer solution. Three technical replicates were performed for each reaction to correct for pipetting errors. The following qPCR profile was used on the LightCycler® instrument (Roche): 3 min of activation phase at 95°C, 40 cycles of 5 s at 95°C, and 30 s at 60°C. Primers are listed in **Table S21**. The *rpl32* gene was used as a reference ^27^ to calculate relative expression levels of *yellow* and EGFP using the manufacturer’s software and the delta-delta Ct method (2^−ΔΔCt^). Differences in the expression of *yellow* and EGFP between the controls and transheterozygous lines were statistically tested using the one-way ANOVA and Tukey’s multiple-comparison test in RStudio (version 1.2.5033).

The RNA samples collected for crosses between the Ub:CasRx-NLS and gRNA lines with relative controls were also used to perform RNA-seq analyses to further validate the qPCR analysis as well as detect the collateral activity of CasRx resulting in misregulation of the whole transcriptome. For these analyses, mRNA was isolated from ~1 μg of total RNA using NEBNext Poly(A) mRNA Magnetic Isolation Module (NEB #E7490). RNA integrity was assessed using the RNA 6000 Pico Kit for Bioanalyzer (Agilent Technologies 5067-1513), and RNA-seq libraries were constructed using NEBNext Ultra II RNA Library Prep Kit for Illumina (NEB #E7770) following manufacturer’s instructions. Libraries were quantified using a Qubit dsDNA HS Kit (ThermoFisher Scientific #Q32854), and the size distribution was confirmed using a High Sensitivity DNA Kit for Bioanalyzer (Agilent Technologies #5067-4626). Libraries were sequenced on an Illumina HiSeq2500 in single read mode with a read length of 50 nt and sequencing depth of 20 million reads per library. Basecalling was performed with RTA 1.18.64 followed by conversion to FASTQ with bcl2fastq 1.8.4.

### Quantification and differential expression analysis

For these studies, the RNA was treated and sequenced as above. The reads were mapped to *Ae. aegypti* genome AaegL5.0 (GCF_002204515.2) supplemented with the PUb-dCas9 transgene sequence using STAR ^28^. Gene expression was then quantified using featureCounts against the NCBI *Ae. aegypti* Annotation Release 101 (GCF_002204515.2_AaegL5.0_genomic.gtf). TPM values were calculated from counts produced by featureCounts and were combined (**Table S4**, **S6**). Hierarchical clustering and PCA analyses were performed in R and were plotted using R package ggplot2. Differential expression analyses between the controls (Ub:CasRx-NLS; U6b-gRNA^*yellow-A*^ and U6b-gRNA^*EGFP*^) and transheterozygous lines (Ub:CasRx-NLS/U6b-gRNA^*yellow*-^ *A* and Ub:CasRx-NLS/U6b-gRNA^*EGFP*^) were performed with DESeq2 (**Table S7**, **S8**, **S11**, **S12**, **S14**, **S15**, **S16**). The Illumina RNA sequencing data have been deposited to the NCBI Sequence Read Archive (SRA), accession number #PRJNA912231 (reviewer link: https://dataview.ncbi.nlm.nih.gov/object/PRJNA912231?reviewer=7ll0k7o7hauq2eh8kbv16mjbj9).

### Phenotypic Screening

To collect eggs laid from single-pair mating events, female and male mosquitoes were allowed to mate for 3 days post-eclosion. Females were given a blood meal for 2 consecutive days. The following day, blood-fed females were independently placed into separate plastic drosophila vials lined with wet paper and plugged with a foam plug. The females were kept in the vials for 2–3 days for egg laying. Following oviposition onto the paper, females were released into a small cage, and the laid eggs were collected, counted, and allowed to mature to full development (~ 4 days) in their original vials. Mature eggs were hatched in their original vial under vacuum overnight. After hatching, the egg papers were removed from the vials to provide more space for larval growth.

At the L3 stage, the progeny were screened, scored, and counted for the expression of opie-2-dsRed, 3xP3-tdTomato, and Opie2-EGFP using a fluorescence-equipped stereoscope (Leica M165FC). The difference in numbers between the total larval counts compared to the total egg counts were considered to have died during embryonic or early larval stages. Surviving transheterozygous individuals were collected for further observation and analysis. Mosquito larvae and pupae were also imaged using a Leica DMC4500 color camera. The hatching rate was calculated as the ratio of the number of larvae to the number of eggs, and the penetrance of the EGFP-targeting phenotype was calculated as the ratio between the number of transheterozygotes showing visual EGFP reduction to the total number of transheterozygotes. Statistical analyses were performed using the one-way ANOVA and Tukey’s multiple-comparison test in RStudio (version 1.2.5033, © 2009–2019).

### Design and selection of the viral target sites

For CHIKV gRNA arrays, CHIKV genomes (469) were downloaded from Virus Pathogen Database and Analysis Resource (ViPR, www.viprbrc.org) and aligned with the CHIKV LR2006_OPY1 strain (GenBank DQ443544) MUSCLE in MegAlign Pro 17 (**Files S1**). Aligned strains were found to have over 90% identity to CHIKV LR2006_OPY1. Alignment of the laboratory strain used (isolate 06113879, Mauritius strain, GenBank MH229986) found 99.64% identity between both strains. Consensus sequences from alignments were then used to identify conserved regions of interest of at least 30nt in length in the non-structural protein (nsp) coding regions of CHIKV. Two suitable target sequences were found in nsp1, nsp2 regions, and two within the nsp4 coding region (**Fig. S5**). We then generated four gRNAs, one gRNA for each of the identified conserved regions (**File S1**, **Table S20**).

### SENSR Assays

SENSR assays were performed as described previously ^19^ using a two-step nucleic acid detection protocol. Briefly, target sequences were first amplified in a 30 min isothermal preamplification reaction by combining reverse transcription with recombinase polymerase amplification (RT-RPA). RT-RPA primers were designed to amplify 30 bp gRNA spacer complement regions flanked by 30 bp priming regions from the synthetic vRNA template while simultaneously incorporating a T7 promoter sequence and “GGG” into the 5′ end of the dsDNA gene fragment to increase transcription efficiency ^29^. The RT-RPA primer sequences and synthetic target sequences can be found in **Table S21**. The RT-RPA was performed at 42°C for 30 min by combining M-MuLV-RT (NEB #M0253L) with TwistAmp Basic (TwistDx #TABAS03KIT). The final conditions for the optimized (28.5% sample input) RT-RPA protocol in a 10-μL reaction are as follows: 5.9 μL rehydration buffer (TwistDx), 0.35 μL primer mix (10 μM each), 0.4 μL reverse transcriptase (200 U/μL), 0.5 μL MgOAc (280 mM), and 2.85 μL vRNA.

The RT-RPA reaction was then transferred to a second reaction, termed the Cas cleavage reaction (CCR), which contained a T7 polymerase and the CasRx ribonucleoprotein. In the second reaction, *in vitro* transcription was coupled with the cleavage assay and a fluorescence readout using 6-carboxyfluorescein (6-FAM). A 6-nt poly-U probe (FRU) conjugated to a 5′ 6-FAM and a 3′ IABlkFQ (Iowa Black Fluorescence Quencher) was designed and custom-ordered from IDT. Detection experiments were performed in 20-μL reactions by adding the following reagents to the 10-μL RT-RPA preamplification reaction: 2.82 μL water, 0.4 μL HEPES pH 7.2 (1 M), 0.18 μL MgCl2 (1 M), 1 μL rNTPs (25 mM each), 2 μL CasRx (55.4 ng/μL), 1 μL RNase inhibitor (40 U/μL), 0.6 μL T7 Polymerase (50 U/μL), 1 μL gRNA (10 ng/μL), and 1 μL FRU probe (2 μM). Experiments were immediately run on a LightCycler 96 (Roche #05815916001) at 37°C for 60 min with the following acquisition protocol: 5 sec acquisition followed by 5 sec incubation for the first 15 min, followed by 5 sec acquisition and 55 sec incubation for up to 45 min. Fluorescence readouts were analyzed by calculating the background-subtracted fluorescence of each timepoint and by subtracting the initial fluorescence value from the final value. Statistical significance was calculated using a one-way ANOVA followed by specified multiple comparison tests (n = 3). The collateral cleavage speed of each gRNA was calculated from the half-maximum fluorescence (HMF) ^19^.

### Vector competence assay

All experiments were performed under biosafety level 3 (BSL-3) conditions in the insectary at the Australian Centre for Disease Preparedness as previously described ^5,30^. The CHIKV (isolate 06113879, Mauritius strain, GenBank MH229986) viral strains grown in Vero cells were used for viral challenge experiments in mosquitoes. Briefly, female mosquitoes were challenged with an infected blood meal (~1 × 10^6^ median tissue culture infective dose [TCID_50_] /mL) through membrane feeding using chicken blood and skin. For determining viral dissemination and infection frequency, whole mosquitoes were collected at 14 days post-challenge, with all deceased mosquitoes collected daily. Daily observations were performed to track mosquito survival during the course of the experiment. The Mantel-Cox statistical test for survival was performed using GraphPad Prism (Version 9.5.0)

### Total RNA extraction and qRT-PCR of viral gene expression

Total RNA was extracted from homogenized whole mosquitoes using the Qiagen RNeasy plus mini kit (Qiagen 74136). Following extraction, cDNA synthesis was performed with ~1 μg of RNA using the Superscript III First-Strand synthesis system (Thermo Fisher 18080051). qRT-PCR was performed with TB Green Premix Ex Taq II (Takara RR820L). Each 20-μl reaction contained 2 μL of cDNA and 10 μL of TB Green solution with a final primer concentration of 200 nM. Three technical replicates were performed for each reaction to correct for pipetting errors. The following qPCR profile was used on the QuantStudio 6 Real-Time PCR system (Thermo Fisher): 30 s at 95°C (hold stage), 40 cycles of 5 s at 95°C, and 30 s at 60°C followed by a melt curve stage of 15 s at 95°C, 60 s at 60°C, and 15 s at 95°C. The *rps17* gene ^27^ was used as a reference to calculate the expression of CHIKV E1 using the delta-delta Ct method (2^−ΔΔCt^). The Mann-Whitney U test was performed using GraphPad Prism (version 9.5.0).

### Ethical conduct of research

All animals were handled in accordance with the Guide for the Care and Use of Laboratory Animals as recommended by the National Institutes of Health and approved by the UCSD Institutional Animal Care and Use Committee (IACUC, Animal Use Protocol #S17187) and UCSD Biological Use Authorization (BUA #R2401).

## Acknowledgements

We thank Judy Ishikawa for mosquito husbandry assistance. This work was supported by funding from a DARPA Safe Genes Program Grant (HR0011-17-2-0047), and NIH awards (R01AI151004, DP2AI152071, and R21AI149161) awarded to O.S.A. We acknowledge the capabilities of the Australian Centre for Disease Preparedness (grid.413322.5) in undertaking this research, including infrastructure funded by the National Collaborative Research Infrastructure Strategy. Figures were created with BioRender.com.

## Data availability

All plasmids and annotated DNA sequence maps are available at www.addgene.com under accession numbers: 191374 (OA-1050T), 191375 (OA-1085F), 191376 (OA-1085L), 194001 (OA-1163A), 194003 (OA-1093B). Raw sequencing data are available at NCBI Sequence Read Archive (SRA), accession number PRJNA912231 (reviewer link: https://dataview.ncbi.nlm.nih.gov/object/PRJNA912231?reviewer=7ll0k7o7hauq2eh8kbv16mjbj9).

## Authors contributions

O.S.A., P.N.P. and E.D.B. conceived and designed the experiments. E.D.B., H.H.L., R.A.M., A.L.D., D.J.B., M.B., T.Y., M.L., I.A., performed molecular and genetic experiments. I.A. performed the RNA sequencing experiments and analysis. A.L.D., M.D., M.J.K., S.J., K.C., K.B. performed viral infection experiments and analysis. All authors contributed to the writing, analyzed the data, and approved the final manuscript.

## Supplementary Information

**Table S1:** RNA sequencing of Ub:CasRx-NLS lines. TPM value

**Table S2:** List of lines used in this study with qPCR data for Ub-CasRx-NES

**Table S3:** hatching data for Yellow

**Table S4:** RNA sequencing of crosses and parental line for yellow analysis (TPM)

**Table S5:** Hatching data for GFP

**Table S6:** RNA sequencing of crosses and parental line for GFP analysis (TPM)

**Table S7:** Differential Expression analysis Ub:CasRx-NLS Versus U6b:gRNAarray_Yellow

**Table S8:** Differential Expression analysis Ub:CasRx-NLS Versus U6b:gRNAarray_GFP

**Table S9:** List of downregulated gene in parental lines for yellow targeting

**Table S10:**List of downregulated genes in parental lines for GFP targeting

**Table S11:** Differential Expression Ub: CasRx Vs. transhets yellow targeting in egg

**Table S12:** Differential Expression U6b:gRNAarray Vs. transhets yellow targeting in egg

**Table S13:** Gene enrichment analysis for yellow targeting

**Table S14:** Differential Expression Ub: CasRx Vs. transhets yellow targeting in larvae

**Table S15:** Differential Expression Ub: CasRx Vs. transhets GFP targeting

**Table S16:** Differential Expression U6b:gRNAarray Vs. transhets GFP targeting

**Table S17:** Gene enrichment analysis for GFP targeting

**Table S18:** Hatching data for yellow targeting by Ub:AeCasRx-NES

**Table S19:** Hatching data for yellow targeting by Ub:AeCasRx-NES

**Table S20:** CHIKV target sites

**Table S21:** Primer sequences used in this study

**File S1:** Alignment of CHKIV consensus sequences used to identify target regions.

**Fig. S1:**
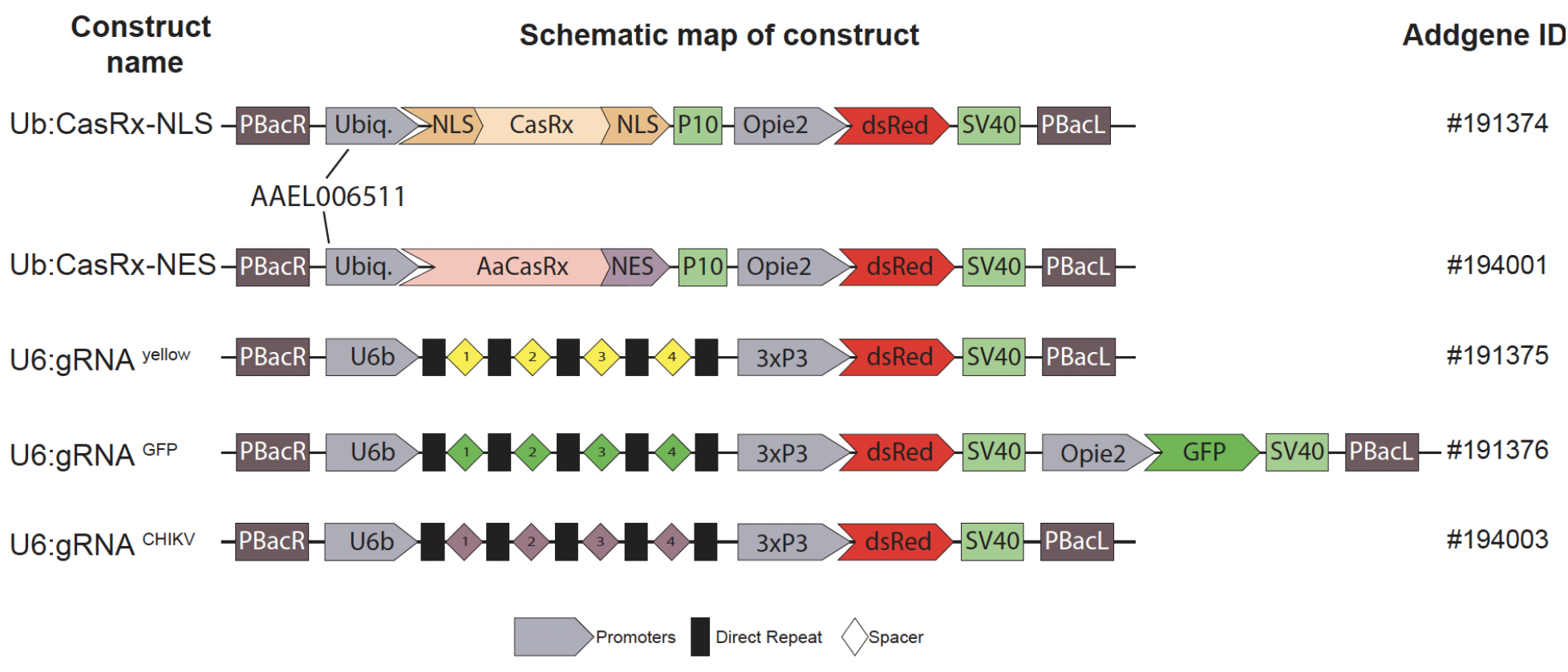
Schematic representation of constructs generated for this study.

**Fig. S2:**
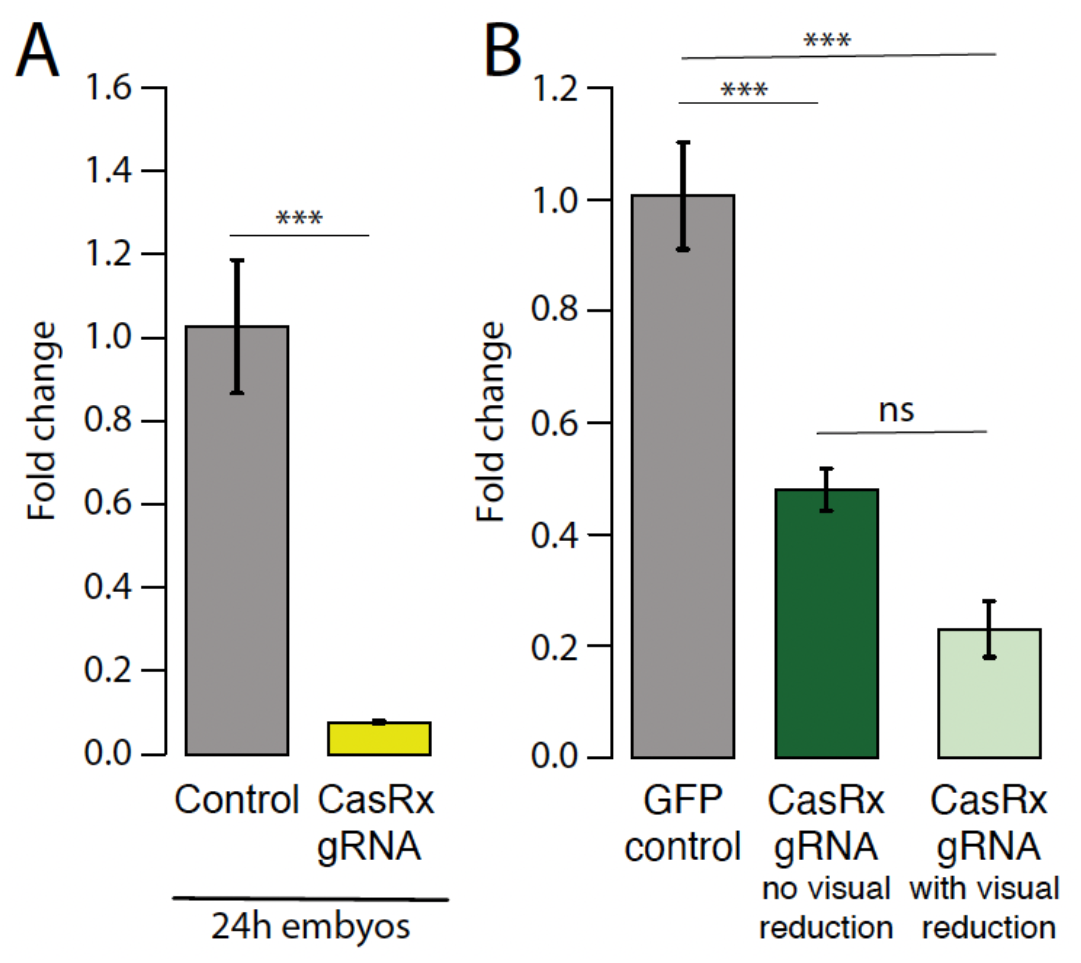
qPCR data validating RNAseq results. **(A)** qPCR result for *yellow* gene in 24–hr-old embryos and surviving larvae. **(B)** GFP expression comparison in controls and transheterozygote larvae with and without visual reduction in GFP. Asterisks indicate significant reduction in hatching rate in transheterozygotes by one-way ANOVA with Tukey’s multiple-comparison test (\*\*\**P* < 0.001).

**Fig. S3:**
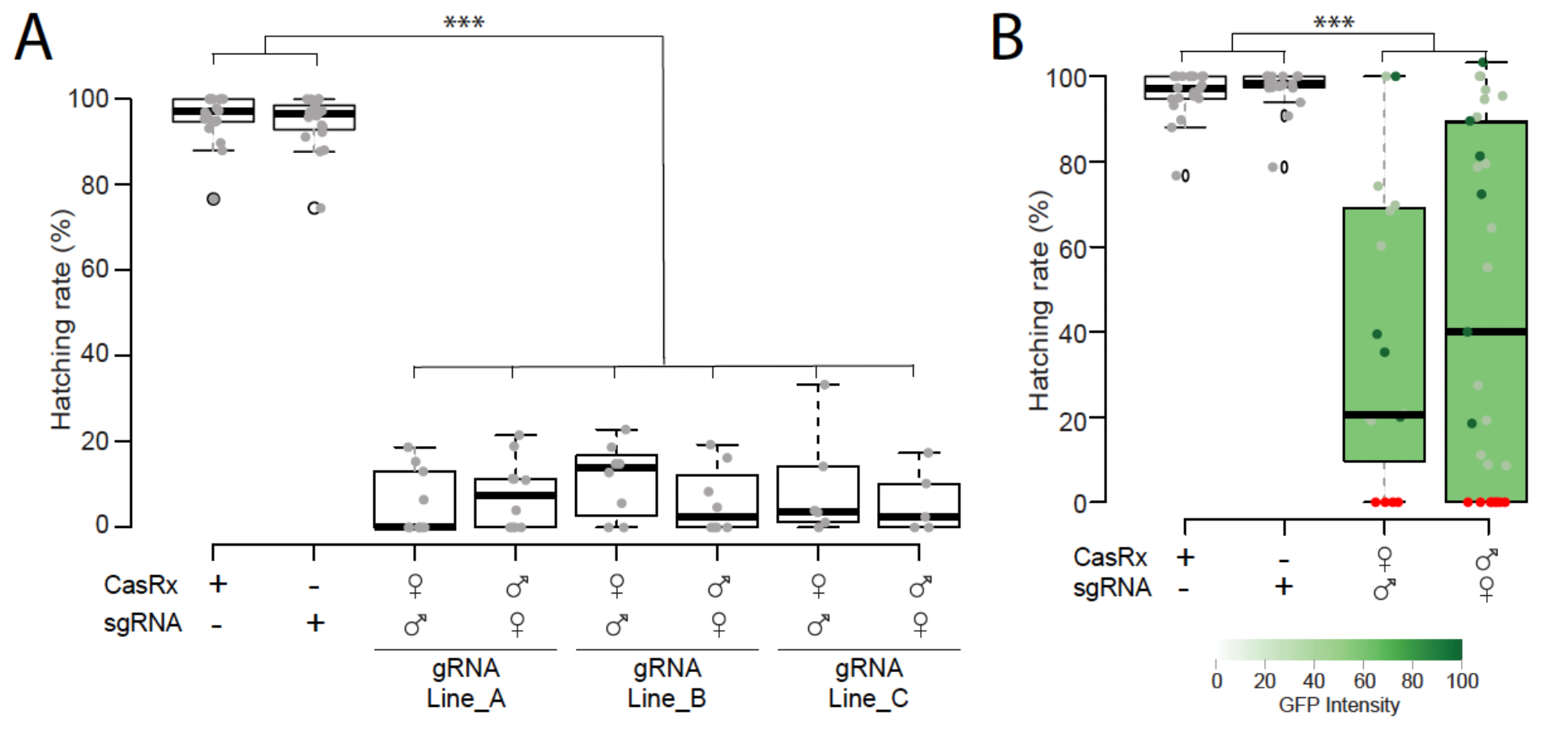
Hatching rate from bidirectional crosses. **(A)** when targeting *yellow* gene and **(B)** when targeting GFP transgene. Phenotype penetrance is depicted by green shading in the box plot, with colors ranging from light green (low EGFP levels) to dark green (high EGFP levels). Red dots represent dead individuals and thus non quantifiable EGFP. Asterisks indicate significant reduction in hatching rate in transheterozygotes by one-way ANOVA with Tukey’s multiple-comparison test (\*\*\**P* < 0.001).

**Fig. S4:**
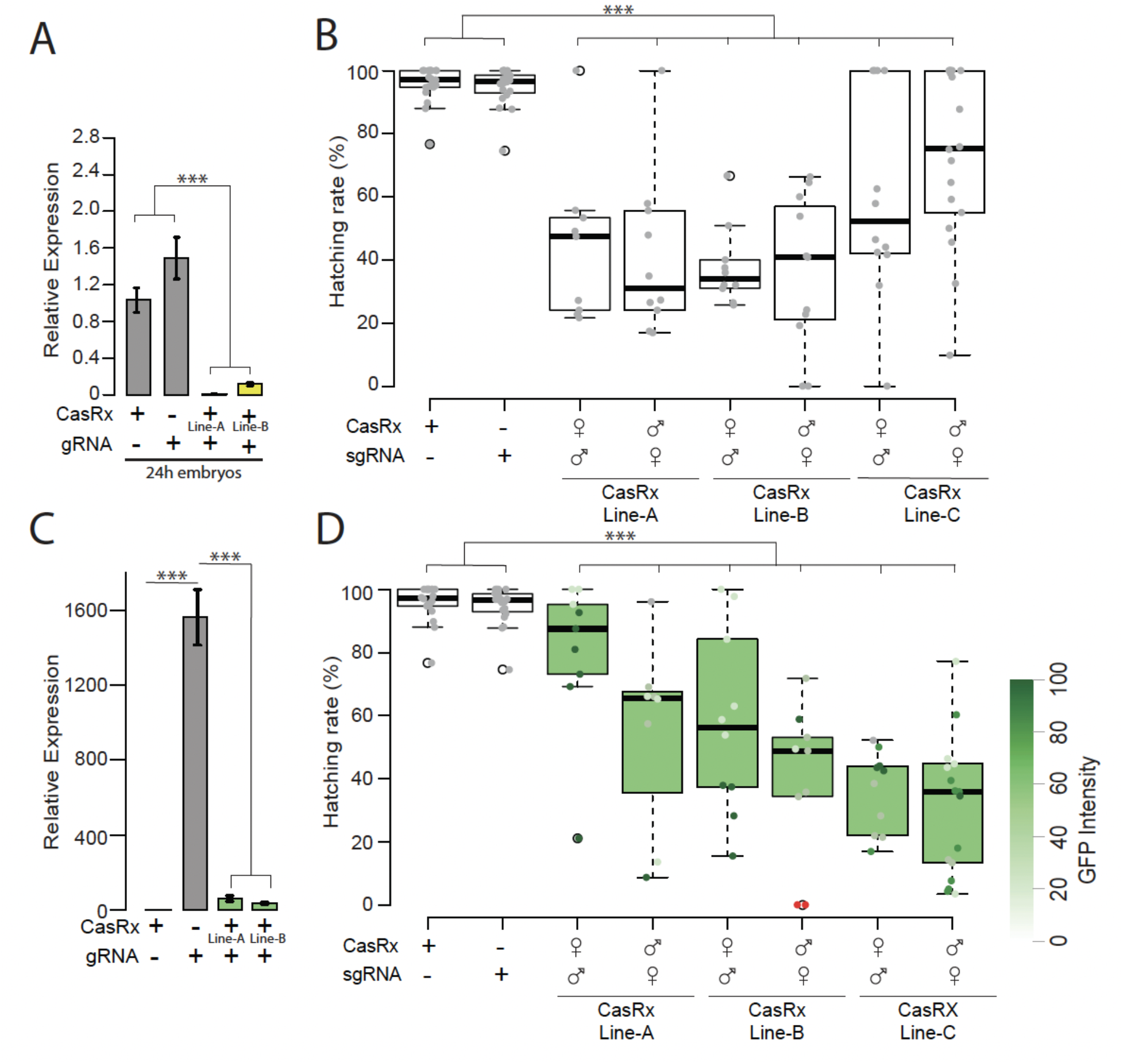
Assessment of yellow and GFP transcript reduction mediated by Ub:AeCasRx-NES. **(A)** qPCR result for *yellow* gene in 24–hr-old embryos, **(B)** Hatching rate from bidirectional crosses using the three Ub:CasRx-NES lines A, B and C (**table S2**). **(C)** qPCR result for *GFP* gene in surviving larvae showing visual reduction in GFP expression. **(D)** Hatching rate from bidirectional crosses using the three Ub:CasRx-NES lines. Phenotype penetrance is depicted by green shading in the box plot, with colors ranging from light green (low EGFP levels) to dark green (high EGFP levels). Red dots represent dead individuals and thus non quantifiable EGFP Asterisks indicate significant differences by one-way ANOVA with Tukey’s multiple-comparison test (*** *P* < 0.001).

**Fig. S5:**
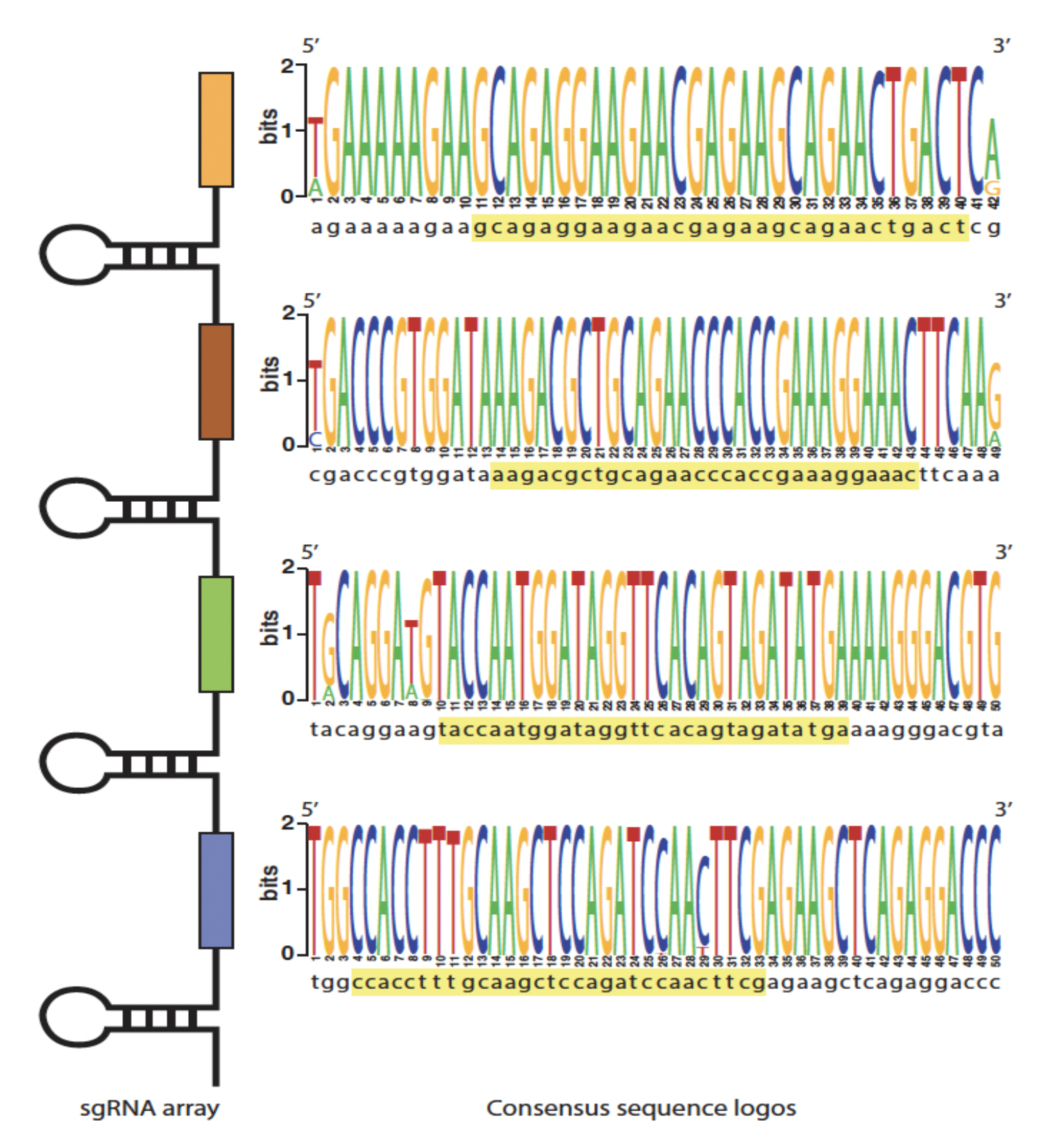
CHIKV consensus sequences of the target sites. Consensus sequences were generated from conserved CHIKV genome regions aligned to the lab strain. They contain the 30nt target region (yellow highlight) used to design the gRNA employed in the study.

**Fig. S6:**
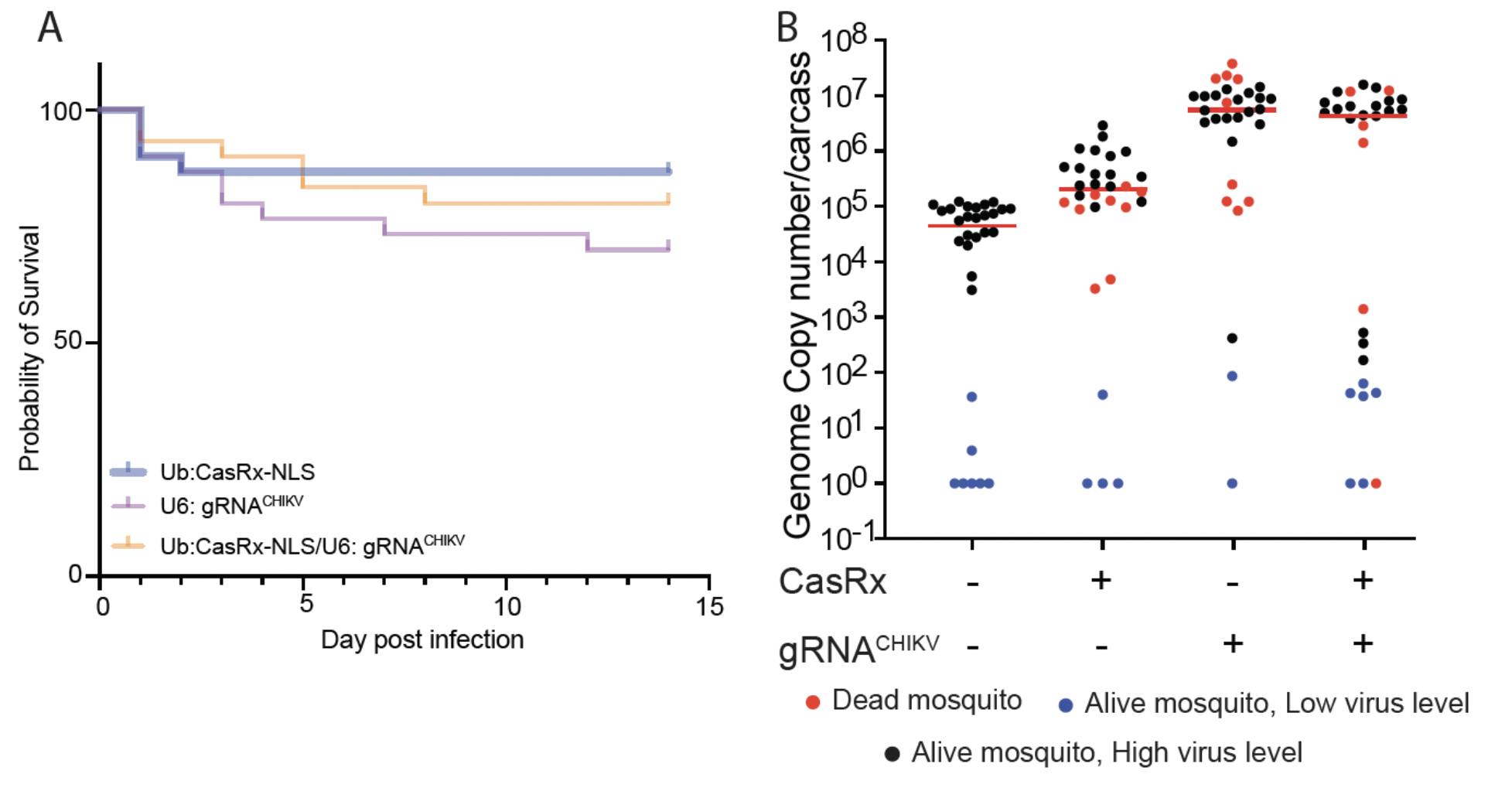
Virus challenge assays using Ub:CasRx-NLS. **(A)** Adult survival curves (log-rank Mantel-Cox test) of CHIKV exposed females per treatment. No significant differences were found between Survival curves. (**B**) The viral genome copy number and infection prevalence of CHIKV were measured after an infected blood meal challenge (n = 30). qRT-PCR was used to assess genome copy number and infection prevalence in individual mosquitoes, with each dot representing the viral load from individual mosquitoes. Each pie-chart indicates the percentage of mosquitos that died by day 5 (in red), `that are alive but with lower virus level of ≤10`^2^ (blue), or that have an high virus level (black). Horizontal red lines indicate the median of the viral loads. Considering the non-normal distribution of viral titers, the median was used to describe central tendency. The non-parametric Mann-Whitney test was used to compare median viral titers, and Fisher’s exact test was used to compare infection prevalence. \**P* < 0.05, \*\**P* < 0.01, \*\*\*\**P* < 0.0001.

